# Non-serotype 2 isolates from healthy pigs are a potential zoonotic reservoir of *Streptococcus suis* genetic diversity and antimicrobial resistance

**DOI:** 10.1101/2021.06.17.447897

**Authors:** Nattinee Kittiwan, Jessica K. Calland, Evangelos Mourkas, Matthew D. Hitchings, Susan Murray, Pakpoom Tadee, Pacharaporn Tadee, Kwanjit Duangsonk, Guillaume Méric, Samuel K. Sheppard, Prapas Patchanee, Ben Pascoe

**Affiliations:** Department of Food Animal Clinics, Faculty of Veterinary Medicine, Chiang Mai University, Chiang Mai, 50100, Thailand; Integrative Research Centre for Veterinary Preventive Medicine, Faculty of Veterinary Medicine, Chiang Mai University, Chiang Mai, 50100, Thailand; Veterinary Research and development center (Upper Northern region), Hang Chat, Lampang, 52190, Thailand; The Milner Centre for Evolution, Department of Biology and Biochemistry, University of Bath, UK; Swansea University Medical School, Swansea University, Singleton Park, Swansea, UK; Faculty of Animal Science and Technology, Maejo University, Chiang Mai, 50290, Thailand; Department of Microbiology, Faculty of Medicine, Chiang Mai University, Chiang Mai, 50200, Thailand; Faculty of Allied Medical Science, Chiang Mai University, Chiang Mai, 50200, Thailand; Department of Zoology, University of Oxford, South Parks Road, Oxford, OX1 3PS, UK

**Keywords:** *Streptococcus suis*, antimicrobial resistance, zoonosis, horizontal gene transfer, mobile elements, one health, gene pool transmission, meningitis

## Abstract

*Streptococcus suis* is a leading cause of bacterial meningitis in SE Asia, with frequent zoonotic transfer to humans associated with close contact with pigs. A small number of invasive lineages are responsible for endemic infection in the swine industry causing considerable global economic losses. A lack of surveillance and a rising trend in clinical treatment failure has raised concerns of growing antimicrobial resistance (AMR) among invasive *S. suis*. The source-sink dynamics between healthy and disease isolates is poorly understood and, in this study, we sample and sequence a collection of isolates predominantly from healthy pigs in Chiang Mai province, Northern Thailand. Pangenome comparisons with a selection of invasive serotype 2 isolates identified increased genetic diversity and more frequent AMR carriage in isolates from healthy pigs. Multiple antimicrobial resistance genes were identified conferring resistance to aminoglycosides, lincosamides, tetracycline and macrolides. All isolates were non-susceptinle to three or more different antimicrobial classes, and 75% of non-serotype 2 isolates were non-susceptible to 6 or more classes (compared to 37.5% of serotype 2 isolates). Antimicrobial resistance genes were found on integrative and conjugative elements (ICE) previously observed in other species, suggesting mobile gene pool which can be accessed by invasive disease isolates.

**Significance statement:** The zoonotic pathogen *Streptococcus suis* causes respiratory disease in pigs and is among the most common causative agents of human clinical bacterial meningitis in SE Asia. We collected isolates from farmed healthy pigs in Northern Thailand, representing a source population from which invasive isolates have recently emerged – linked to the pork production industry. Pangenome characterisation of the isolates revealed a reservoir of genetic diversity and antimicrobial resistance suggesting that One Health approaches may be beneficial in tackling the increase in antimicrobial resistance.

## Background

More than half the world’s pork meat is produced in SE Asia, and China alone is home to nearly half the world’s live pigs. The United Nations’ Food and Agricultural Organization estimated that China produces around half of the billion pigs reared worldwide (Gilbert et al., 2018). This massive increase in agricultural intensification has brought significant challenges in animal welfare, including infection control. Among the most common infections to Asian herds is a respiratory disease caused by *Streptococcus suis* (VanderWaal and Deen, 2018). Infection with *S. suis* occurs mainly in piglets and growing pigs and can lead to septicemia with sudden death, arthritis, endocarditis, meningitis (Dutkiewicz et al., 2017; Gottschalk and Segura, 2019; Segura, 2020). *S. suis* infections accounted for a loss of over US$11 million to the pork industry in Thailand alone in 2019 (Rayanakorn et al., 2020). This expanded niche for *S. suis* has provided opportunities for zoonotic infection, which are frequently reported worldwide following increased exposure to pigs, often in farm workers, slaughterhouse workers, and butchers (Goyette-Desjardins et al., 2014; van Samkar et al., 2015; Dutkiewicz et al., 2017). However in SE Asia, particularly in Northern Thailand, where there is a tradition of consuming raw pork dishes, *S. suis* infection is one of the most common causative agents of clinical bacterial meningitis (Takeuchi et al., 2017; Rayanakorn et al., 2019).

Human zoonotic *S. suis* infections predominantly arise from a single virulent lineage, thought to have emerged in the 1920s alongside the intensification of the pork production industry. However, no consistent genomic differences between pig and human disease isolates have been observed (Weinert et al., 2015). This may be related to the fact that isolates from healthy (asymptomatic) pigs have not been well studied but it is known that disease-associated isolates have fewer genes overall but more that encode putative virulence factor (Weinert et al., 2019; Murray et al., 2021). Serotyping of the *S. suis* capsular polysaccharides is often used in epidemiological studies, with 29 *S. suis sensu stricto* serotypes described to date (Athey et al., 2016a; Segura et al., 2016). *S. suis* serotype 2 is the most virulent and is frequently isolated from diseased pigs and human clinical cases (Hughes et al., 2009; Okura et al., 2016); however, non-serotype 2 isolates (often isolated from healthy pigs) represent an extensive reservoir of genetic diversity (Zhang et al., 2011; Baig et al., 2015; Okura et al., 2019; Stevens et al., 2019).

Widespread use of antimicrobial drugs in the pig production industry has driven an increase in antimicrobial resistance (AMR) (Van Boeckel et al., 2015; WHO, 2017). Imprudent use of colistin in pork production as a growth enhancer (since the 1970s) encouraged the development of resistance in *E. coli* (and other gram-negative bacteria), which has diminished the effectiveness of antibiotics used in human medicine (Liu et al., 2016; Delannoy et al., 2017; Patchanee et al., 2020a). Furthermore, there is a rising trend in multi-drug resistant (MDR) zoonotic pathogens, such as Salmonella, that pose a significant public health threat (Prasertsee et al., 2019; Patchanee et al., 2020b; Tadee et al., 2021). Regulation of veterinary use of antibiotics is difficult in low- and middle-income countries, which consequently have some of the highest AMR levels (Nguyen et al., 2016). For example, in Thailand alone, infections with antimicrobial-resistant bacteria are estimated to cause up to 38,000 human deaths each year (Pumart P, 2012).

A lack of surveillance and rise in clinical treatment failure has raised concerns of growing antimicrobial resistance among invasive *S. suis* (Pathanasophon et al., 2013). Furthermore, the source-sink dynamics among commensal and disease-causing isolates are poorly understood. In this study, we sample and sequence a collection of isolates predominantly from healthy pigs in Chiang Mai province, Northern Thailand. Pangenome comparisons with a selection of invasive serotype 2 isolates identified increased genetic diversity and more frequent AMR carriage in isolates from healthy pigs. Antimicrobial resistance genes were found on integrative and conjugative elements (ICE) previously observed in other species, suggesting mobile gene pool which can be accessed by invasive disease-causing isolates.

## Results

All *S. suis* samples collected from healthy pigs in Chiang Mai province, Thailand were identified by PCR (**Table S1**) as non-serotype 2 isolates. From the 138 isolates we collected, 25 were randomly selected for whole genome sequencing. An additional 11 isolates from lab archives, previously collected from Chiang Mai were added to the dataset to include representative isolates from pig disease and invasive human infection. In total, the dataset used consisted of 36 isolates, of which 8 isolates (22.2%) were serotype 2, including isolates from human clinical cases (n=2), diseased pigs (n=2), and healthy pigs (n=4) and 28 isolates (77.8%) of non-serotype 2 *S. suis* from healthy pigs (**Figure 1A; Table S2**).

**Figure 1.**
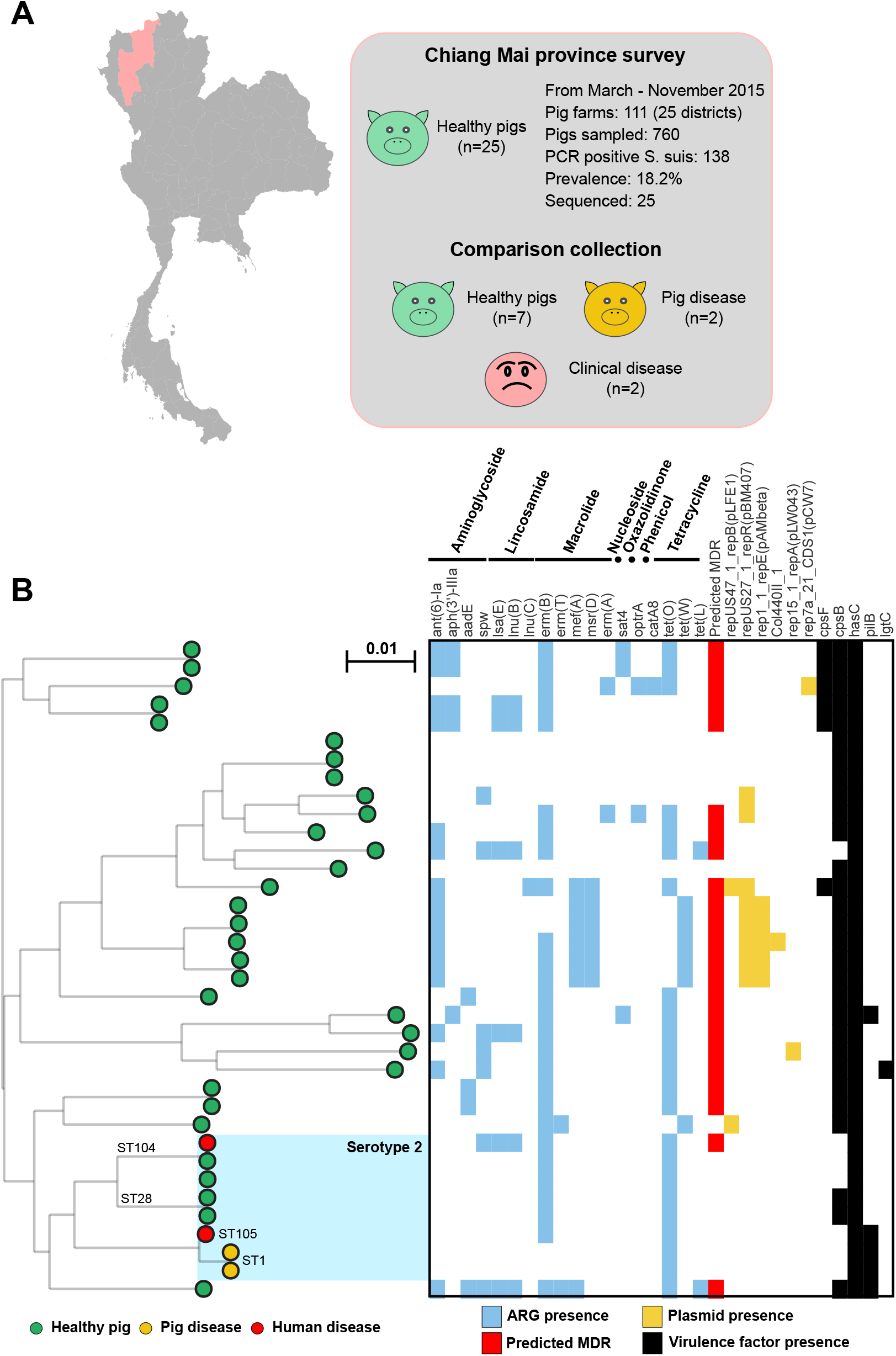
**A:** Isolates were collected as part of a survey of healthy pigs in Chiang Mai province, Thailand. **B:** Population structure of selected sequenced isolates compared with other serotype 2 genomes from the same region. All core genes (present in ≥95% of isolates) from the reference genome (1,348 genes) were used to build a gene-by-gene alignment (n = 36; 1,202,840 bp). A maximum-likelihood phylogeny was constructed with IQ-TREE, using a GTR model and ultrafast bootstrapping (1,000 bootstraps; version 1.6.8; Nguyen et al; 2015; Hoang et al 2018). Scale bar represents genetic distance of 0.01. Leaves are colored by disease status and host: samples from healthy pigs are green; diseased pigs are yellow; and samples from human clinical cases are red. Serotype 2 isolates are shaded in blue, with common STs annotated. The presence of antimicrobial resistance genes, known plasmids and virulence genes identified using ABRICATE and NCBI, PlasmidFinder and VfDb databases are indicated by coloured blocks. Interactive visualization is available on Microreact: https://microreact.org/project/Ssuis-ns2 (Argimon *et al*; 2016).

### Non-serotype 2 isolates are a reservoir of antimicrobial resistance

No non-serotype 2 isolates were responsible for disease in either pigs or humans. A maximum-likelihood phylogeny constructed from a concatenated gene-by-gene core genome alignment (1,348 genes) revealed a highly structured population (**Figure 1B; Supplementary file S1**). Serotype 2 isolates clustered together, including the previously described sequence types ST-1, ST-28, ST-104 and ST-105. Pairwise average nucleotide identity (ANI) comparisons suggested that non-serotype 2 isolates (75.1% identical) were more diverse than serotype 2 isolates (98.1% identical) in the core genome (**Figure 2AB**). This was supported by (pairwise) clustering of the core and accessory genome using PopPUNK (Lees et al., 2019), which identified divergence in the accessory genomes of the serotype 2 isolates (**Figure 2C**). Together the pangenome of all 36 isolates comprised 5,004 gene clusters, with 1,348 core genes present in at least 95% of isolates representing ~27% of the pangenome; or ~68% of the average *S. suis* genome (1,993 ORFs in BM407; **Figure 2D; Table S3**). Typically, invasive serotype 2 isolates have smaller genomes but contain more virulence-related genes (Weinert et al., 2015). In our dataset, this was also true with serotype 2 isolates having smaller genomes on average (**Table S2**), and the virulence associated *pilB* gene was found in 75% (n=3 of 4) of invasive isolates, but only 7% of isolates from healthy pigs (n=2 of 28) (**Figure 1B; Table S4**).

**Figure 2.**
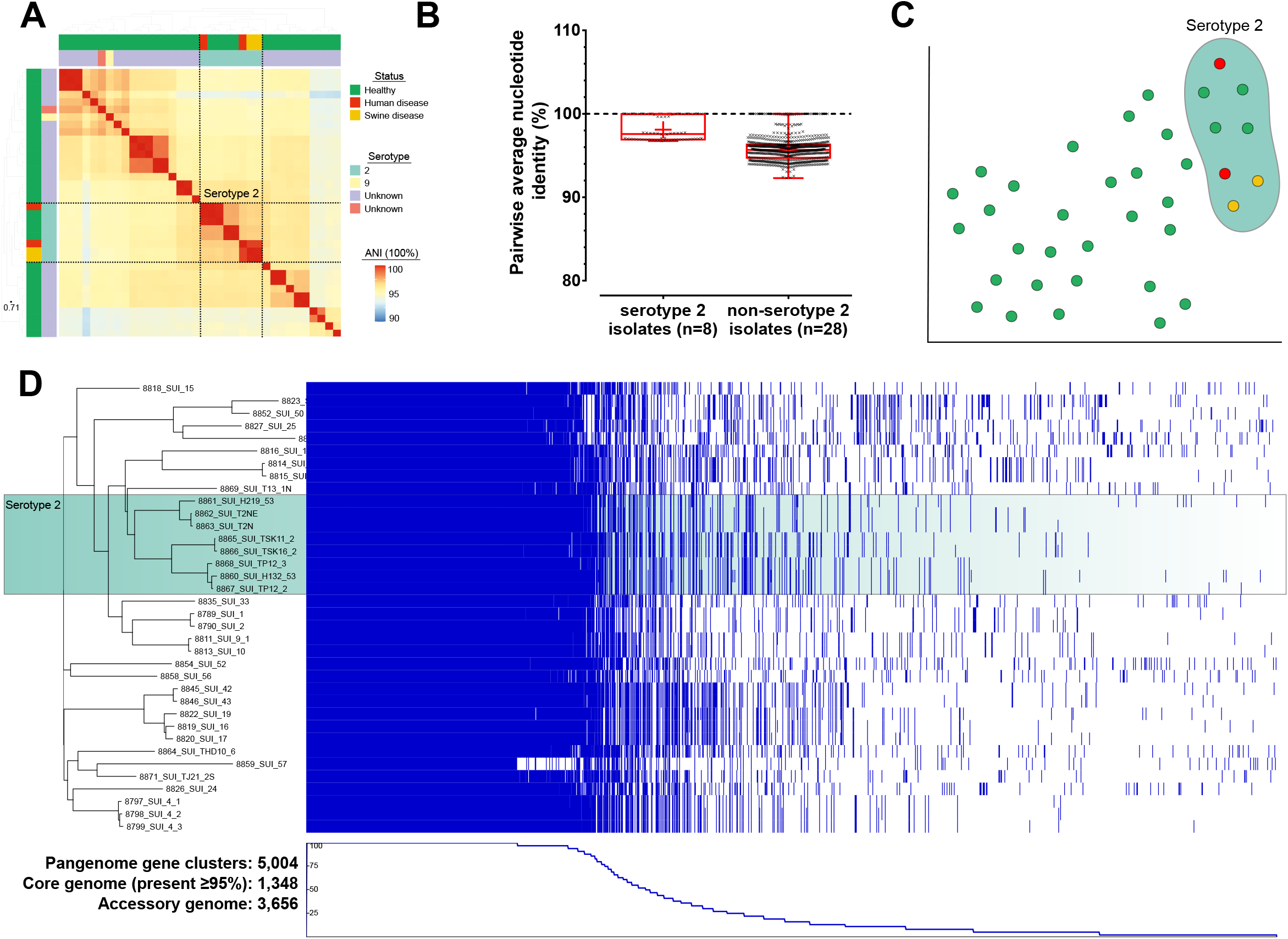
**A:** Heatmap of pairwise average nucleotide identity (ANI). Highly similar pairwise comparisons are colored in red to blue for the most dissimilar isolates. The cluster of serotype 2 isolates are boxed. **B**: Summary of pairwise comparisons between serotype 2 and non-serotype 2 isolates. **C**: PopPUNK pairwise accessory distances visualized with t-SNE clustering in microreact: https://microreact.org/project/Ssuis-ns2 (Argimon *et al*; 2016). **D**: Visualisation of the pangenome (PIRATE) with phandango, including estimation of the core (gene present in 95% or more isolates) and accessory genome composition (Bayliss et al; 2019; Hadfield et al 2018).

### Widespread AMR determinants in S. suis isolates from healthy pigs

We scanned all 36 genomes for known determinants of antimicrobial resistance (AMR) through nucleotide comparisons (≥70% sequence identity) with the NCBI database (Coordinators, 2018; Bortolaia et al., 2020) and identified 18 resistance genes from 7 different antimicrobial classes (**Figure 1B; Table S5**). Loci conferring putative resistance to aminoglycosides (*aadE, ant*(*6*)-*la, aph*(*3*’)-*III* and *spw*); macrolides (*erm*(*A*), *erm*(*B*), *erm*(*T*), *mef*(*A*) and *msr*(*D*)); lincosamides (*lsa*(*E*), *lnu*(*B*) and *lnu*(*C*)); tetracycline (*tet*(*w*), *tet*(*L*) and *tet*(*O*)); oxazolidinone (*optrA*); nucleoside (*sat4*) and chloramphenicol (*catA8*) were found in 32 isolates (89%). On average, fewer antibiotic resistance genes were identified in the serotype 2 isolates (5 genes) compared to non-serotype 2 isolates (18 genes; **Table S5**). All 18 of the resistance genes were detected in the non-serotype 2 isolates from healthy pigs, but only five of the potential AMR genes *spw*, *lsa*(*E*), *erm*(*B*), (*lnu*(*B*), and *tet*(*O*) were found in the eight serotype 2 isolates, including *tet*(*O*) which was present in all serotype 2 isolates. At least one antimicrobial resistance gene from three or more antimicrobial classes was found in 21 out of 36 isolates (58%), and only one out of these was a *S. suis* serotype 2 isolate.

### Evidence of mobility of AMR genes among S. suis from healthy pigs

Comparison of nucleotide sequences from all the genomes with the PlasmidFinder database (Carattoli et al., 2014) identified loci identified on six putative integrative and conjugative elements (ICE), including pLFE1, pBM407, pAMbeta, Col440II, pLW043, and pCW7 (**Figure 1B; Table S6**). All putative ICE elements were identified in non-serotype 2 isolates (39%; 11 of 28). Two of these ICE elements have previously been characterized in invasive *S. suis*, pBM407 (accession: FM252033) and pAMbeta (accession: AE002565.1). The pBM407 plasmid described in *S. suis* contained AMR genes conferring resistance to tetracycline (*tetO, tetL*), chloramphenicol (acetyltransferase), erythromycin (*ermB*) and a dihydrofolate reductase (Holden et al., 2009). However, plasmids from two different isolates with variation in gene content hint at an underlying diversity - and this potential composite architecture was evidenced by differences in the AMR gene complement (Holden et al., 2009). All serotype 2 isolates contain the *tetO* locus, and 75% (6 of 8) contain the *ermB* locus which are described as members of the pBM407 ICE element, but no other pBM407 genes are identified by this method (**Table S6**). Additional plasmids not previously described in *S. suis* were also identified using MOB-suite, which compares genome sequences with all described plasmids in the NCBI database (**Table S7**; (Robertson and Nash, 2018; Robertson et al., 2020).

### Widespread antimicrobial resistance in non-serotype 2 isolates

Disk diffusion assays were used to determine antimicrobial susceptibility of the isolates to 18 antimicrobial agents, from 9 antimicrobial categories. Most isolates were highly susceptible to linezolid (100%; n=36), amoxicillin-clavulanic acid (97%; n=35), ceftiofur (94%; n=34), amoxicillin (83%; n=30) and ampicillin (81%; n=29). High levels of resistance were observed against lincomycin (100%; n=36), clindamycin (97%; n=35), tetracycline (92%; n=33), doxycycline (92%; n=33), kanamycin (89%; n=32), oxytetracycline (83%; n=30), erythromycin (69%; n=25) and gentamycin (31%; n=11) (**Table 1**). There was a statistically significant difference in antimicrobial susceptibility between *S. suis* serotype 2 and non-serotype 2 isolates for gentamycin (*p*-value: 0.037), chloramphenicol (*p*-value: 0.010), and penicillin G (*p*-value: 0.001; Pearson’s Chi-square test and Fisher’s exact test) (**Table 1**). Multi-drug resistance (MDR) is defined as an isolate that is non-susceptible to at least one antimicrobial agent from three different antimicrobial categories (Sweeney et al., 2018). In this study, we will consider all non-susceptible isolates as resistant. All 36 *S. suis* isolates were resistant to three or more antibiotic classes (**Figure 3A; Table 1**). Most (87.5%; 7 of 8) serotype 2 isolates were resistant to four and five antimicrobial categories; while three quarters (21 of 28) of non-serotype 2 isolates were resistant to 6, 7 and 8 antimicrobial categories (46.4, 25 and 3.6%, respectively; **Figure 3B**).

**Table 1.**
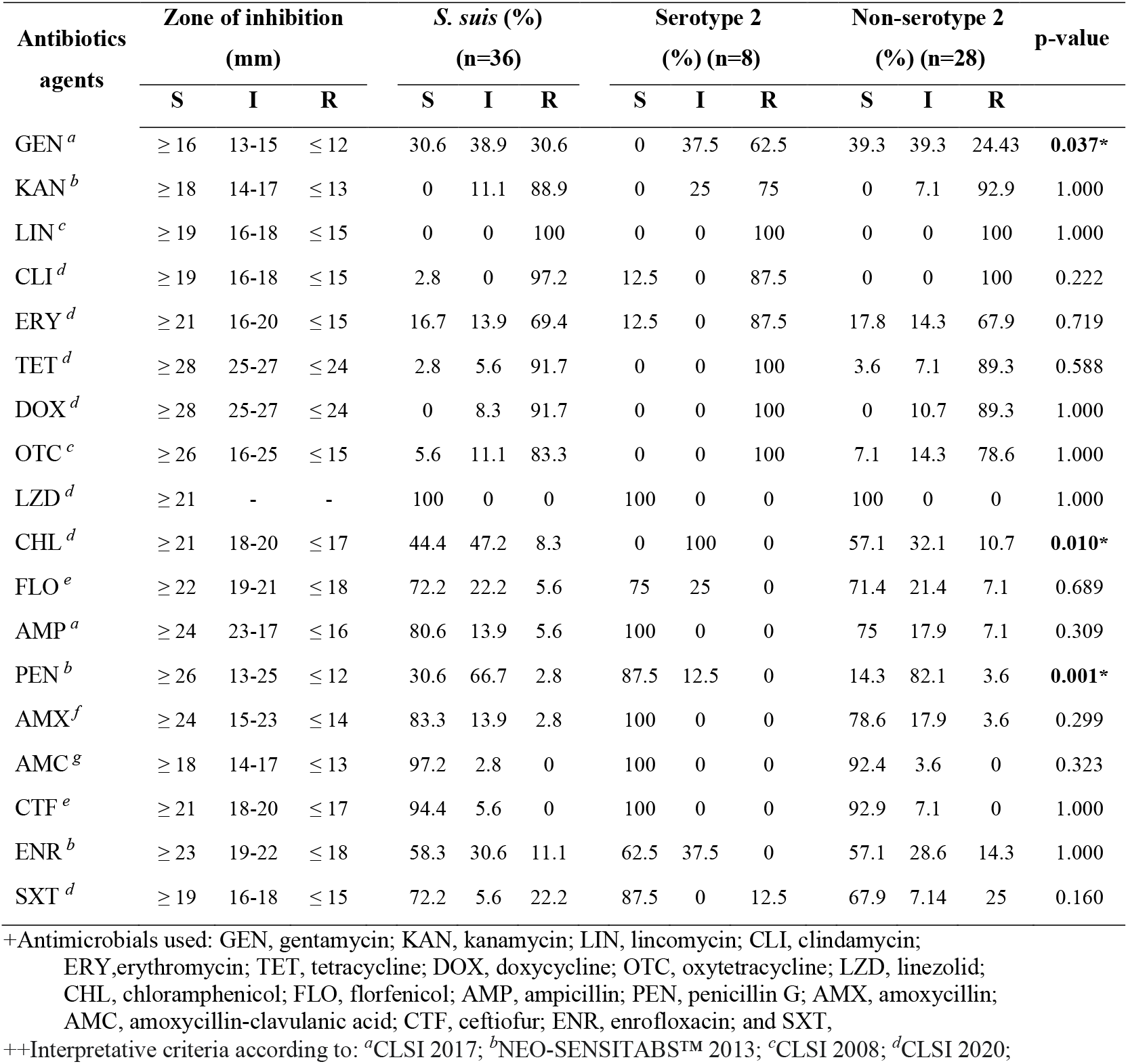
Antimicrobial susceptibility test results by disk diffusion method of 36 *S. suis*, grouped by serotype. Susceptible (S), intermediate (I) and resistant (R) phenotypes are indicated. Asterisk (*) indicates statistical significance by Pearson’s Chi-square test and Fisher’s exact test, *p*-value < 0.05.

**Figure 3.**
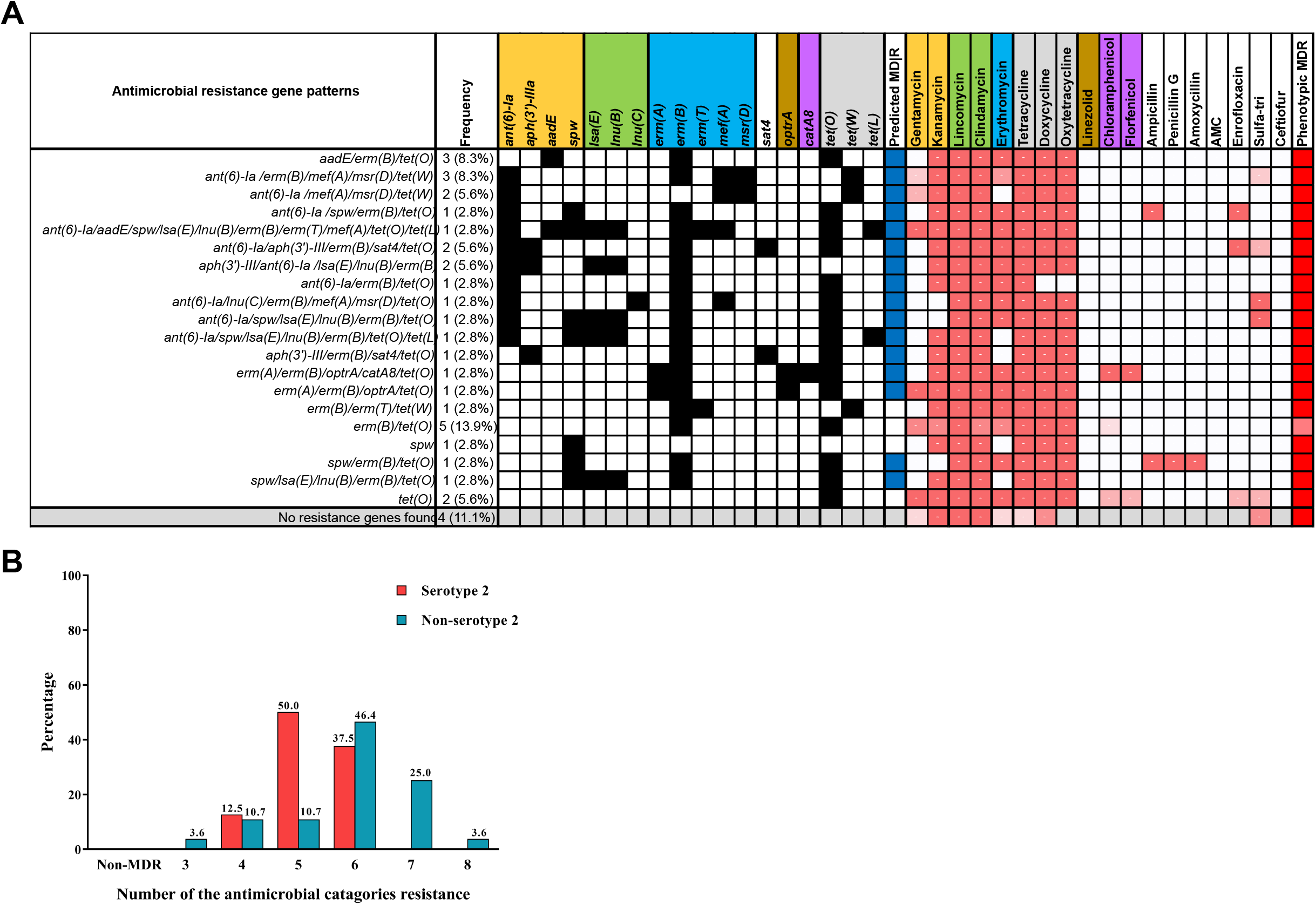
**A**: Distribution of antimicrobial resistance patterns (black blocks indicate antimicrobial resistance genes identified in the genomes) summarized alongside their corresponding phenotypic resistance outcomes (red blocks indicated phenotypic resistance). Predicted (blue) and phenotypic (red) MDR is also indicated. **B:** Summary of the number of different antimicrobial classes to which each isolate demonstrated phenotypic resistance. Isolates resistant to three or more different antimicrobial classes were characterized as MDR.

### Diverse AMR profiles in S. suis isolates from healthy pigs

Increased diversity in the core, accessory, and plasmid content of non-serotype 2 isolates was associated with increased AMR conferred by 21 different antimicrobial resistance gene (ARG) profiles (A–U; **Figure 3A; Table *2***). The most common ARG pattern included the *erm*(*B*) and *tet*(*O*) genes (13.9%), followed by *aadE/erm*(*B*)/*tet*(*O*) and *ant*(*6*)-*Ia* /*erm*(*B*)/*mef*(*A*)/*msr*(*D*)/*tet*(*W*) gene patterns (8.3%). All putative resistance genes were absent in 4 non-serotype 2 strains (11.1%). Non-serotype 2 isolates demonstrated greater variation in ARG content, with only three resistance gene patterns (B, C and P) found exclusively in serotype 2 isolates.

**Table 2.** Antimicrobial resistance gene patterns of 36 *S. suis* isolates.

| **Pattern** | **Antimicrobial resistance gene patterns** | **Strains** | **Frequency** |
| --- | --- | --- | --- |
| **A** | *spw* | FH57 | 1 (2.8%) |
| **B** | *tet(O)* | DP-T2NE <sup>†, β</sup>, DP-T2N <sup>†, β</sup> | 2 (5.6%) |
| **C** | *erm(B)/tet(O)* | H219-53<sup>α,β</sup>, TSK11-2<sup>β</sup>, TSK16-2<sup>β</sup>, TP12-2<sup>β</sup>, TP12-3<sup>β</sup> | 5 (13.9%) |
| **D** | *aadE/erm(B)/tet(O)* | FH11, FH12, FH52 | 3 (8.3%) |
| **E** | *ant(6)-Ia/erm(B)/tet(O)* | THD10-6 | 1 (2.8%) |
| **F** | *erm(B)/erm(T)/tet(W)* | FH13 | 1 (2.8%) |
| **G** | *spw/erm(B)/tet(O)* | FH25 | 1 (2.8%) |
| **H** | *aph(3')-III/erm(B)/sat4/tet(O)* | FH20 | 1 (2.8%) |
| **I** | *ant(6)-Ia /spw/erm(B)/tet(O)* | FH51 | 1 (2.8%) |
| **J** | *ant(6)-Ia /mef(A)/msr(D)/tet(W)* | FH16, FH17 | 2 (5.6%) |
| **K** | *erm(A)/erm(B)/optrA/tet(O)* | TJ21-2S | 1 (2.8%) |
| **L** | *aph(3')-III/ant(6)-Ia /erm(B)/sat4/tet(O)* | FH9, FH10 | 2 (5.6%) |
| **M** | *aph(3')-III/ant(6)-Ia /lsa(E)/lnu(B)/erm(B)* | FH1, FH2 | 2 (5.6%) |
| **N** | *ant(6)-Ia /erm(B)/mef(A)/msr(D)/tet(W)* | FH19, FH42, FH43 | 3 (8.3%) |
| **O** | *erm(A)/erm(B)/optrA/catA8/tet(O)* | FH33 | 1 (2.8%) |
| **P** | *spw/lsa(E)/lnu(B)/erm(B)/tet(O)* | H132-53 <sup>α, β</sup> | 1 (2.8%) |
| **Q** | *ant(6)-Ia/lnu(C)/erm(B)/mef(A)/mrd(D)/tet(O)* | FH24 | 1 (2.8%) |
| **R** | *ant(6)-Ia/lsa(E)/lnu(B)/erm(B)/tet(O)/tet(L)* | FH15* | 1 (2.8%) |
| **S** | *ant(6)-Ia/spw/lsa(E)/lnu(B)/erm(B)/tet(O)* | FH50 | 1 (2.8%) |
| **T** | *ant(6)-Ia/aadE/spw/lsa(E)/lnu(B)/erm(B)/erm(T)/mef(A)/tet(O)/tet(L)* | T13-1N | 1 (2.8%) |
| **U** | No resistance genes found | FH4-1, FH4-2, FH4-3, FH56 | 4 (11.1%) |
| **Total** |  |  | **36 (100%)** |

\* = *S. suis* serotype 9, <sup>α</sup> = *S. suis* from the human case, <sup>β</sup> = *S. suis* serotype 2, <sup>†</sup> = *S. suis* from diseased pigs

### Phenotypic and genotypic concordance in antimicrobial resistance

When we compared phenotypic non susceptibility (zones of inhibition) with the presence of specific ARGs, often there was no clear correlation (**Table 3; Figure S1**). There was little correlation between gentamycin, kanamycin, lincosamide, clindamycin resistance and the presence of any specific ARG. Widespread resistance to tetracycline (97.2% non susceptible; **Table 1**), doxycycline and oxytetracycline correlated with the presence of ARGs, including *tet*(*O*), *tet*(*W*) and *tet*(*L*) (OR > 1; **Table 3**). The strongest link between phenotype and ARG was observed for erythromycin resistance and the presence of *erm*(*B*) (OR= 32.5, *p* =0.002; **Table 3**). Phenotypic resistance to erythromycin correlated well with the ARGs *erm*(*A*), *erm*(*T*), *mef*(*A*) and *msr*(*D*); and 72.2% of erythromycin resistant isolates contained the *erm*(*B*) gene (**Table 3; Figure S1**). There were not enough resistant isolates to properly assess the correlation between the presence of genes linked to resistance to chloramphenicol (n=1) and florfenicol (n=1) and none of the isolates were resistant to the first generation oxazolidinones, linezolid, despite identification of the corresponding *optrA* resistance gene (**Table 3; Figure S1**).

**Table 3.** Concordance of antimicrobial resistance phenotype and genotypes. Presence of resistance genes (G+) and number of phenotypically non-susceptible isolates (P+) indicated. Asterisk indicates *p*-value < 0.05 by Pearson’s chi-square test, and Yate’s correction for continuity.

| Antimicrobial agents | AMR genes | Characterization of phenotypic and genotypic resistance (n=36) |  |  |  | Concordance of phenotypic and genotypic resistance |  |
| --- | --- | --- | --- | --- | --- | --- | --- |
|  |  | P+/G+ | P-/G+ | P+/G- | P-/G- | OR (95% CI) | p-value |
| GEN | *ant(6)-Ia* | 8 (22.2%) | 7 (19.4%) | 16 (44.4%) | 5 (13.9%) | 0.36 (0.08- 1.42) | 0.175 |
|  | *aph(3')-III* | - | 5 (13.89%) | 24 (66.7%) | 7 (17.9%) | 0.06 (0.005-0.46) | 0.02 |
|  | *aadE* | 3 (8.3%) | 1 (2.8%) | 21 (58.3%) | 11 (30.6%) | 1.57 (0.21-22.1) | 0.851 |
|  | *spw* | 3 (8.3%) | 3 (8.3%) | 21 (58.3%) | 9 (25%) | 0.43 (0.09-2.17) | 0.635 |
| KAN | *ant(6)-Ia* | 15 (41.7%) | - | 21 (58.3%) | - | 0.71 (0.04-14.4) | 0.615 |
|  | *aph(3')-III* | 5 (13.9%) | - | 31 (86.1%) | - | 0.16 (0.01-3.64) | 0.714 |
|  | *aadE* | 4 (11.1%) | - | 32 (88.9%) | - | 0.13 (0.01-2.94) | 0.611 |
|  | *spw* | 6 (16.7%) | - | 30 (83.3%) | - | 0.2 (0.01-4.4) | 0.805 |
| LIN, CLI | *lsa(E)* | 6 (16.7%) | - | 30 (83.3%) | - | 0.2 (0.01-4.4) | 0.805 |
|  | *lnu(B)* | 6 (16.7%) | - | 30 (83.3%) | - | 0.2 (0.01-4.4) | 0.805 |
|  | *lnu(C)* | 1 (2.8%) | - | 35 (97.2%) | - | 0.03 (0.002-1.06) | 0.199 |
| ERY | *erm(A)* | 2 (5.6%) | - | 28 (77.8%) | 6 (16.7%) | 0.43 (0.04-7.16) | 0.745 |
|  | *erm(B)* | 26 (72.2%) | 1 (2.8%) | 4 (11.1%) | 5 (13.9%) | 32.5 (3.79-390) | **0.002\*** |
|  | *erm(T)* | 2 (5.6%) | - | 28 (77.8%) | 6 (16.7%) | 0.43 (0.04-7.16) | 0.917 |
|  | *mef(A)* | 6 (16.7%) | 1 (2.8%) | 24 (66.7%) | 5 (13.9%) | 1.25 (0.16-16.9) | 0.706 |
|  | *msr(D)* | 5 (13.9%) | 1 (2.8%) | 25 (69.4%) | 5 (13.9%) | 1 (0.12-13.9) | 0.548 |
| TET | *tet(O)* | 23 (63.9%) | - | 12 (33.3%) | 1 (2.3%) | 1.92 (0.09-37.8) | 0.758 |
|  | *tet(W)* | 6 (16.7%) | - | 29 (80.6%) | 1 (2.8%) | 0.207 (0.01-4.55) | 0.821 |
|  | *tet(L)* | 2 (5.6%) | - | 33 (91.7%) | 1 (2.8%) | 0.06 (0.003-1.7) | 0.368 |
| DOX | *tet(O)* | 23 (63.9%) | - | 13 (36.1%) | - | 1.77 (0.09-34.9) | 0.721 |
|  | *tet(W)* | 6 (16.7%) | - | 30 (83.3%) | - | 0.2 (0.01-4.40) | 0.805 |
|  | *tet(L)* | 2 (5.6%) | - | 3 (94.4%) | - | 0.06 (0.03-1.65) | 0.357 |
| OTC | *tet(O)* | 23 (63.9%) | - | 11 (30.6%) | 2 (5.6%) | 4.18 (0.43-62.7) | 0.573 |
|  | *tet(W)* | 6 (16.7%) | - | 28 (77.8%) | 2 (5.6%) | 0.43 (0.04-7.16) | 0.917 |
|  | *tet(L)* | 2 (5.6%) | - | 32 (88.9%) | 2 (5.6%) | 0.12 (0.01-2.65) | 0.571 |
| LZD | *optrA* | - | 2 (5.6%) | - | 34 (94.4%) | 17 (0.61-327) | 0.357 |
| CHL | *catA8* | 1 (2.8%) | - | 19 (52.8%) | 16 (44.4%) | 0.84 (0.04-16.9) | 0.541 |
| FLO | *catA8* | 1 (2.8%) | - | 9 (25%) | 26 (72.2%) | 2.89 (0.14-56.6) | 0.947 |

## Discussion

*S. suis* were cultured and identified from 18.6% of pigs swabbed in this study (138 of 760 samples), which is within the range previously reported for the prevalence in farmed pigs and slaughterhouses in the same area of Thailand (Padungtod et al., 2010; Kongkaew et al., 2012). This level of prevalence was significantly lower than the *S. suis* prevalence previously reported in pigs from other provinces in Northern Thailand, such as Lampang (64.8%) and Phayao (61.4%) (Pathanasophon et al., 2013). These and other studies in Northern Thailand reported high prevalence of serotype 2 (5.6-43%) and serotype 7 (8.2-14.3%) isolates (Padungtod et al., 2010; Thongkamkoon et al., 2017). However, we mainly identified non-serotype 2 isolates, with only a single isolate typed as serotype 9 and no serotype 2 isolates identified during this survey. This variation is likely due to differences in sampling, as we prioritised collection from healthy pigs. Invasive disease isolates have shown biogeographical variation, and with competition and serotype replacement noted among virulent *S. suis* serotypes (Flores et al., 1993; Hughes et al., 2009; Hadjirin et al., 2020). Serotype 9 is most common in diseased pigs from Europe and has a low pathogenic potential in humans. However, a rare case of serotype 9 infection in humans has recently been reported in Thailand (Goyette-Desjardins et al., 2014; Kerdsin et al., 2017).

Difficulties in (capsule) serotyping *S. suis* (*sensu lato*) isolates, where previously typed *S. suis* isolates are now designated as other *Streptococcsu* species hint at an ambiguous species designation and variation in non-serotype 2 isolates (Prüfer et al., 2019; Hatrongjit et al., 2020). This is further supported by characterisation of divergent *S. suis* isolates by whole genome sequencing (Baig et al., 2015). The extent to which these represent stable lineages is unclear, with the rate of lineage turnover in *S. suis* seldom investigated (Calland et al., 2020; Hadjirin et al., 2020). We identified increased variation in the core and accessory genomes of non-serotype 2 isolates (**Figures 1 and 2**). Serotype 2 isolates are typically found to have smaller genomes than non-invasive isolates (Baig et al., 2015; Weinert et al., 2015) and our collection of mostly non-invasive non-serotype 2 isolates were consistently larger (with more genes; **Table S2**) than the *S. suis* reference genome (BM407: 2,170,810 bp) and selected invasive isolates (**Table S2**). Despite smaller genomes, these invasive isolates also tend to carry more virulence-related genes and all serotype 2 isolates in our collection carried the *pilB* gene, which is associated with the brain cell invasion required to cause meningitis in humans and pigs (Maisey et al., 2007). It has been suggested that this reduction in genome size may be due to gene loss, including core metabolism genes for nutrients that can be scavenged from the host; and a streamlining of functional/redundant elements (Weinert and Welch, 2017; Murray et al., 2021).

A plug-and-play theory of bacterial accessory genomes (Young, 2016; McInerney et al., 2017; Sheppard et al., 2018), where diversity in bacterial phenotypes can be conferred by a mobile pool of genes that are readily gained and lost, enables the acquisition of rapid adaptive genomic changes that can be spread through the population via recombination (Redondo-Salvo et al., 2020). Host switching and zoonotic infection complicate analyses of source and sink dynamics and attribution of AMR elements (Dearlove et al., 2016; Mourkas et al., 2019). Here, we focus primarily on the potential reservoir of infection and characterize variation in the gene pool from which invasive disease isolates have arisen. Where resistance is conferred by a single (or few) nucleotide substitution(s), it is impossible to tell from sequence data if HGT or point mutation were responsible (Zhao et al., 2016; Bortolaia et al., 2020). For other classes of antibiotics, the literature provides clear evidence for HGT of genes (Florez-Cuadrado et al., 2017; Wang et al., 2018; Patchanee et al., 2020a; Redondo-Salvo et al., 2020). For example, the pBM407 plasmid characterised in the pBM407 *S. suis* reference genome mobilizes *tetO*, *tetL, emrB, cat* and *dfr* genes between isolates (Holden et al., 2009; Hoa et al., 2011). Including these putative tetracycline resistance genes, our analyses identified 18 accessory genes associated with resistance to 7 antimicrobial classes.

We identified genes with described roles in resistance to aminoglycosides, macrolides, lincosamides, tetracycline, nucleoside, oxazolidinones, and phenicols. Most isolates were predicted to be MDR (80.6%). The most common antimicrobial resistant genes identified were associated with resistance to macrolides and tetracycline. More than 80% of isolates contained at least one gene predicted to confer macrolide resistance (Palmieri et al., 2011). The presence of *erm*(*B*) and *mef*(*A*) genes are consistent with previous studies, where *erm*(*B*) is strongly linked with macrolide-lincosamide-streptogramin B (MLS_B_) resistance and presented in 59-90% of macrolide-resistant *S. suis* isolates from pigs (Martel et al., 2001; Zhang et al., 2015; Tan et al., 2020). The resistant gene, *erm*(*T*) has been detected in *S. agalactiae, S. pyogenes*, and other erythromycin-resistant isolates of group D *Streptococci* (Chen et al., 2012; Zhang et al., 2015; Yongkiettrakul et al., 2019), our identification of *erm*(*T*) in this study suggests potential within-genus HGT.

The most common tetracycline resistance gene detected was *tet*(*O*) in over half of the isolates (63.9%) (**Table 3**). An alternative ribosomal protein, *tet*(*M*) is also often associated with tetracycline resistance in *S. suis* (Palmieri et al., 2011; Bojarska et al., 2016) but was not observed among our isolates. In addition, we detected tet(*L*) and *tet*(*W*) genes, which have not often been reported in *S. suis*, among non-serotype 2 isolates from healthy pigs. Corresponding phenotypic resistance to tetracycline was reported in over 90% of isolates, which is consistent with global data reporting widespread resistance to tetracycline and macrolides, likely related to the prophylactic use in agriculture (Soares et al., 2014; Yongkiettrakul et al., 2019; Mourkas et al., 2020; Tan et al., 2020). AMR may play a role in increasing numbers of treatment failures (Hughes et al., 2009; Gurung et al., 2015; Yongkiettrakul et al., 2019), and in our study, despite widespread MDR, we observed phenotypic susceptibility to all three of the recommended antimicrobials used to treat clinical *S. suis* meningitis (penicillin, ceftiofur, and ceftriaxone) (van Samkar et al., 2015; Seitz et al., 2016). However, some β-lactam resistant strains (18-27%) were found in the non-clinical strain of *S.suis* (Soares et al., 2014; Yongkiettrakul et al., 2019; Segura et al., 2020). Despite this, β-lactam usage in pig production should be closely monitored, especially where there is prophylactic use in healthy pigs.

Concordance between the presence of predicted ARGs and phenotypic resistance was poor for most antimicrobial, and we report widespread phenotypic resistance, even in the absence of a predictive resistance element (**Figure 3**). Given the enhanced genetic diversity and lack of clear characterization of this disease reservoir, it is possible that additional resistance elements have yet to be fully described. A recent study by Hadjirin et al. identified more than 20 novel *S. suis* AMR determinants (Hadjirin et al., 2021). Even in the absence of direct antimicrobial selective pressure, broad spectrum use of antibiotics act on all bacterial species in the microbiome; and this bystander effect can confer resistance on bacterial species that are not the target of the antimicrobial treatment (Tedijanto et al., 2018; Morley et al., 2019). Enrofloxacin is widely used to treat other types of bacterial infection in the respiratory and digestive systems of livestock animals, and in our collection more than 40% of isolates were resistant to this antibiotic (Lakkitjaroen et al., 2011; Yongkiettrakul et al., 2019). Spectinomycin is often used in pig production and other livestock animals combined with lincomycin (Bosman et al., 2019; Wang et al., 2020). Clusters of AMR genes (*aadE-spw-lsa*(*E*)-*lnu*(*B*)) have been identified in *Staphylococci* and *Enterococci* associated with lincosamides resistance (Li et al., 2014; Huang et al., 2016). We identified the combination of spectinomycin and lincosamide resistance in one serotype 2 isolate and 2 non-serotype 2 isolates from healthy pig. Individually, we identified spectinomycin and lincosamides resistance genes in a small number of isolates, as has previously been reported in invasive *S. suis* isolates (Athey et al., 2016b; Bojarska et al., 2016).

The plasmid-borne chloramphenicol resistance gene, *catA8* (McHugh et al., 2020; Yan et al., 2020), and the *optrA* gene that confers transferable combined resistance to oxazolidinones (linezolid) and phenicols (chloramphenicol and florfenicol) (Brenciani et al., 2016; Bender et al., 2018; Zhou et al., 2019) are reported here for the first time in *S. suis*. Although phenotypic susceptibility was recorded to linezolid, the isolates were resistant to chloramphenicol and florfenicol. Recently, this gene has been found in oxazolidinone-resistant *S. suis* isolates in China (Huang et al., 2017; Du et al., 2019; Huang et al., 2019). This is further evidence of the unintended effect of broad-spectrum antimicrobials, such as the oxazolidinones linezolid and tedizolid, which are highly effective against Gram-positive bacteria (Sztanke et al., 2004) but rarely used in the pig production industry. However, florfenicol has been used in livestock animals for therapeutic purposes and there is documented transfer of plasmids carrying *optrA* between different Gram-positive bacteria (Wang et al., 2015). Twenty-one different resistance gene patterns were observed, with *erm*(*B*) and *tet*(*O*) found together in 62.5% (5 of 8) of serotype 2 isolates, as previously observed (Athey et al., 2016b). Most non-serotype 2 isolates possessed ARGs to at least three antimicrobial classes (up to seven; 22/28, 78.6%). Several genetic elements, including ICE carrying antimicrobial resistance genes such as *optrA, ermB, tetM, tetO*, and *tetW* have been reported in *S. suis* (Holden et al., 2009; Athey et al., 2016b), however plasmid elements were found only in non-serotype 2 isolates in this study.

## Conclusion

We collected isolates from 760 healthy pigs reared in the pork industry in Northern Thailand. Through comparison of 36 whole genome sequences, we identified increased genetic diversity in these non-serotype 2 carriage isolates, from which the more invasive and pathogenic serotype 2 isolates emerge. Corresponding diversity was also seen in the breadth and diversity of AMR determinants which conferred increased phenotypic non-susceptibility. This genetically diverse reservoir of *S. suis* pose a public health risk with the potential for transmission to more invasive isolates, broadening their spectrum of antimicrobial resistance. Extensive phenotypic resistance is observed to antimicrobials that are not typically used to treat this infection. This can be partly explained by the co-occurrence of resistance genes on ICEs. However, little phenotypic resistance was observed to β-lactams, which remain the prescribed antimicrobial for *S. suis* infection in Thailand. Continued surveillance and more stringent control of antimicrobial usage within the pork industry will be necessary to monitor a growing AMR threat in *S. Suis*.

## Methods

### Ethical approval

This study was carried out according to the guidelines for the care and use of laboratory animals (National Research Council, 2010). The study protocol was approved by the Faculty of Veterinary Medicine’s Animal Care and Use Committee (Protocol number S24/2559).

#### Sample collection

Samples were collected between March and November 2015, with a total of 760 tonsil swab samples collected from 111 pig farms in 25 districts of Chiang Mai province, Thailand. All swab samples were kept in Stuart transport medium (Oxoid, UK) and transported to the laboratory at 4 °C within 24 hrs of collection.

Live fattening pigs were swabbed, and *S. suis* identified in 138 samples (18.2%). Of 138 *S. suis* isolates obtained from healthy pigs, only one isolate (0.7%) was confirmed as *S. suis* serotype 9. Meanwhile, all the remaining 137 isolates (99.3%) were negative to serotypes serotype ½/1/2/7/9/14 by PCR identification and classified as non-serotype 2 strains. Among 138 strains, 25 strains were randomly selected for WGS. In addition, 11 isolates were selected from laboratory archives (collected as part of another study of farmed pigs in Chiang Mai province during 2015) and sequenced for comparison between serotype 2 and non-serotype 2 isolates.

#### Bacterial identification and growth

Tonsil swab samples were inoculated onto 5% sheep blood agar plates (Oxoid, UK) and incubated at 37°C for 24 hours. *S. suis* isolates were identified by biochemical characterization (Quinn et al., 1994), and small (approximately 1 mm in diameter) transparent alpha-hemolysis and non-hemolysis colonies of Gram-positive cocci with negative catalase test were selected for further screening. Criteria for presumptive identification of *S. suis* included no growth on 6.5% NaCl agar, a negative Voges-Proskauer (VP) test, and production of acid in trehalose, lactose, sucrose, salicin and inulin broths, but no acid production in glycerol, sorbitol, and mannitol. A multiplex polymerase chain reaction (PCR) using primers specific to the 16S rRNA gene was used to confirm the identification of *S. suis* and capsular gene types 1 or 14, 2 or 1/2, 7, and 9, which are the most prevalent serotypes recovered from diseased pigs, as described in **Table S1** (Wisselink et al., 2000; Wisselink et al., 2002; Marois et al., 2004).

#### Antimicrobial susceptibility testing

Antimicrobial susceptibility tests were performed using the disk diffusion method in accordance with the recommendations of the Clinical and Laboratory Standards Institute (CLSI) (CLSI, 2012). Eighteen antibiotic drugs from 9 antibiotic groups were used in the test, including aminoglycoside (gentamycin, 10 μg; and kanamycin, 30 μg), lincosamides (lincomycin, 10 μg; and clindamycin, 2 μg), macrolides (erythromycin, 15 μg), tetracyclines (tetracycline, 30 μg; doxycycline, 30 μg; and oxytetracycline, 30 μg), oxazolidinone (linezolid, 30 μg), phenicols (chloramphenicol, 30 μg; and florfenicol 30 μg), beta-lactams (ampicillin, 10 μg; penicillin G, 10 units; amoxycillin, 10 μg; amoxycillin-clavulanic acid, 30 μg; and ceftiofur, 30 μg), fluoroquinolones (enrofloxacin, 5 μg) and folate inhibitors (sulfamethoxazole/trimethoprim, 1.25/23.75 μg) (Oxoid. Hamshire, UK). Diameter breakpoints were assessed according to the guidelines described in Table 1 (CLSI, 2002, 2008, 2012; Howe and Andrews, 2012; NEO-SENSITABS™, 2013; CLSI, 2017, 2018, 2020). Pearson’s Chi-square test and Fisher’s exact test were performed using IBM SPSS Statistics version 22 (IBM, New York, USA) to determine the difference of antimicrobial susceptibility between *S. suis* serotype 2 and non-serotype 2. The association between antimicrobial-resistant phenotype and genotype was tested by Pearson’s chi-square test, and Yate’s correction for continuity was applied where required. Statistically significant associations were shown as odds ratios (ORs) with 95% confidence intervals (CI). Results were considered statistically significant when a two-tailed *p*-value ≤ 0.05.

#### Genome sequencing and assembly

Twenty-five *S. suis* isolates from pigs with no clinical signs of *S. suis* infection (healthy pigs) were randomly selected for sequencing from the 138 recovered samples. Our collection was augmented with two archived isolates derived from tissue samples of pigs with clinical signs of *S. suis* infection (diseased pigs) that were submitted to the Veterinary Medicine Research and Development Center (Upper Northern region) of the National Institute of Animal Health (Thailand), and a further nine isolates from the Faculty of Medicine at Chiang Mai University. In total, our collection included 32 healthy pigs, two diseased pigs, and two human clinical samples cultured from the blood of meningitis patients. All strains were cultured in Todd-Hewitt-broth at 37 °C for 18-24 hrs, and genomic DNA was extracted using the QIAamp DNA Minikit (QIAgen^®^). Whole-genome sequencing (WGS) using a multiplex sequencing approach was performed on an Illumina Miseq genome sequencer (Illumina, Cambridge UK) using Nextera XT libraries and third generation MiSeq reagent kits. Paired-end short reads of 300 bp were filtered, trimmed, and assembled *de novo* with SPAdes version 3.7 (Bankevich et al., 2012). The average number of contiguous sequences (contigs) in 36 *S. suis* genomes was 160 for an average total assembled sequence size of 2.22 Mbp. The average N50 contig length (L50) was 66,810 and the average GC content was 41.3%. Short read data are available on the NCBI SRA, associated with BioProject PRJNA418954. Assembled genomes and supplementary material are available from FigShare (10.6084/m9.figshare. 13385465; individual accession numbers and assembled genome statistics in **Table S**2).

#### Population structure and phylogeny

A multisequence alignment was created from concatenated gene sequences of all core genes (found in >95% isolates) from the reference genome, BM407 (Holden et al., 2009) using MAFFT (Katoh et al., 2002) on a gene-by-gene basis (Morley et al., 2019)(size: 1,202,840 bp; **Supplementary file S1**). Maximum-likelihood phylogenies were reconstructed with IQ-TREE (version 1.6.8) using the GTR+F+I+G4 substitution model and ultra-fast bootstrapping (1,000 bootstraps) (Nguyen et al., 2015); and visualized on Microreact (Argimón et al., 2016): https://microreact.org/project/Ssuis-ns2

#### Accessory genome characterization

All unique genes present in at least one isolate (the pangenome) were identified by automated annotation using PROKKA (version 1.13) followed by PIRATE, a pangenomics tool that allows for orthologue gene clustering in bacteria (Seemann, 2014; Bayliss et al., 2019). We defined genes in PIRATE using a wide range of amino acid percentage sequence identity thresholds for Markov Cluster algorithm (MCL) clustering (45, 50, 60, 70, 80, 90, 95, 98). The pan-genome of all 36 isolates contained 5,004 genes, of which 1,348 genes were shared by all isolates (>95%) and defined the core genome (**Table S3**). Pairwise core and accessory genome distances were compared using PopPUNK (version 1.1.4) (Lees et al., 2019), which uses pairwise nucleotide kmer comparisons to distinguish shared sequence and gene content to identify accessory genome divergence in relation to the core genome. A two-component Gaussian mixture model was used to build a network to define clusters (Components: 41; Density: 0.0579; Transitivity: 0.9518; Score: 0.8967).

#### Identification of virulence-associated genes

The accessory genome of each isolate was characterized, including detection of antimicrobial resistance genes, putative virulence factors, and known plasmid genes using ABRICATE (version 0.9.8) (https://github.com/tseemann/abricate) and the NCBI, VfDB, and PlasmidFinder databases (10th September 2019 update; **Tables S4, S5 and S6**) (Carattoli et al., 2014; Coordinators, 2018; Liu et al., 2019; Bortolaia et al., 2020). ABRICATE was used to identify antimicrobial resistance genes in the sequenced genomes by comparison with the NCBI database of 1,726 resistance genes covering 15 antimicrobial agent types; including genes associated with resistance to aminoglycosides, β-lactam, colistin, fluoroquinolone, fosfomycin, fusidic acid, glycopeptide, MLS-macrolide-lincosamide-streptogramin B, nitroimidazole, oxazolidinones, phenicols, rifampicin, sulphonamides, tetracycline, and trimethoprim. An astringent threshold of 98% identity was used for reporting a match between a gene in the NCBI database and the input genome. T-test and Fisher’s exact test assessed statistical significance at 5% significance.

## Declarations

### Data availability

Short read data are available on the NCBI SRA, associated with BioProject PRJNA418954 (https://www.ncbi.nlm.nih.gov/bioproject/PRJNA418954). Assembled genomes and supplementary material are available from FigShare: doi: 10.6084/m9.figshare.13385465.

### Author contributions

NK: acquisition, analysis, and interpretation of data; drafted manuscript.

JKC: analysis, and interpretation of data; revised manuscript.

EM: analysis, and interpretation of data; revised manuscript.

MDH: acquisition and interpretation of data; revised manuscript.

SM: acquisition and interpretation of data; revised manuscript.

PakT: acquisition and interpretation of data; revised manuscript.

PacT: acquisition interpretation of data; revised manuscript.

KD: acquisition and interpretation of data; revised manuscript.

GM: interpretation of data; revised manuscript.

SKS: interpretation of data; revised manuscript.

PP: conceptualized and designed work; acquisition, analysis, and interpretation of data; drafted manuscript.

BP: conceptualized and designed work; acquisition, analysis, and interpretation of data; drafted manuscript.

## Acknowledgments

We are grateful to all the technicians, scientists, and non-technical staff of the Veterinary Research and Development center (Upper northern region), Lampang, Thailand, and Swansea University medical school, Swansea, UK, for their cooperation and support.

## Supplementary material

**Table S1**: Primers used for species and serotype identification.

**Table S2**: Summary of isolate genome statistics

**Table S3**: Summary of core and accessory genome characterization with PIRATE

**Table S4**: Summary of virulence genes identified by comparison with the VfDB database.

**Table S5**: Summary of AMR genes identified by comparison with the NCBI database.

**Table S6**: Summary of plasmid genes identified by comparison with the PlasmidFinder database.

**Table S7**: Matrix of gene presence for all plasmids identified by MOB-suite.

**Figure S1**: The effect of each ARG on phenotypic resistance diffusion diameters for aminoglycosides, lincosamides, tetracyclines, phenicols, oxazolidinone and macrolide.

**Supplementary file 1**: Alignment file.

## List of abbreviations

AMC: Amoxycillin-Clavulanic acid
AMP: Ampicillin
AMR: Antimicrobial Resistance
AMX: Amoxycillin
ARGs: Antimicrobial Resistance Genes
CARD: Comprehensive Antibiotic Resistance Database
CHL: Chloramphenical
CLI: Clindamycin
CTF: Ceftiofur
DOX: Doxycycline
ENR: Enrofloxacin
ERY: Erythromycin
FLO: Florfenicol
GEN: Gentamycin
ICE: Integrative and Conjugative Elements
KAN: Kanamycin
LIN: Lincomycin
LZD: Linezolid
MDR: Multiple Drug Resistance
MLS: Macrolide-Lincosamide-Streptogramin B
NCBI: National Center for Biotechnology Information
OTC: Oxytetracyclin
PEN: Penicillin G
SXT: Sulfamethoxazole/Trimethoprim
TET: Tetracycline
VFDB: Virulence Factor Database

## References

Argimón, S., Abudahab, K., Goater, R.J.E., Fedosejev, A., Bhai, J., Glasner, C. et al. (2016) Microreact: visualizing and sharing data for genomic epidemiology and phylogeography. Microbial genomics 2: 11.

Athey, T.B., Teatero, S., Lacouture, S., Takamatsu, D., Gottschalk, M., and Fittipaldi, N. (2016a) Determining Streptococcus suis serotype from short-read whole-genome sequencing data. BMC Microbiol 16: 162.

Athey, T.B., Teatero, S., Takamatsu, D., Wasserscheid, J., Dewar, K., Gottschalk, M., and Fittipaldi, N. (2016b) Population structure and antimicrobial resistance profiles of Streptococcus suis serotype 2 sequence type 25 strains. PloS one 11: e0150908.

Baig, A., Weinert, L.A., Peters, S.E., Howell, K.J., Chaudhuri, R.R., Wang, J. et al. (2015) Whole genome investigation of a divergent clade of the pathogen *Streptococcus suis*. Frontiers in microbiology 6:1191.

Bankevich, A., Nurk, S., Antipov, D., Gurevich, A.A., Dvorkin, M., Kulikov, A.S. et al. (2012) SPAdes: a new genome assembly algorithm and its applications to single-cell sequencing. Journal of computational biology 19: 455–477.

Bayliss, S.C., Thorpe, H.A., Coyle, N.M., Sheppard, S.K., and Feil, E.J. (2019) PIRATE: A fast and scalable pangenomics toolbox for clustering diverged orthologues in bacteria. GigaScience 8:10, giz119.

Bender, J.K., Cattoir, V., Hegstad, K., Sadowy, E., Coque, T.M., Westh, H. et al. (2018) Update on prevalence and mechanisms of resistance to linezolid, tigecycline and daptomycin in enterococci in Europe: towards a common nomenclature. Drug Resistance Updates 40: 25–39.

Bojarska, A., Molska, E., Janas, K., Skoczyńska, A., Stefaniuk, E., Hryniewicz, W., and Sadowy, E. (2016) *Streptococcus suis* in invasive human infections in Poland: clonality and determinants of virulence and antimicrobial resistance. European Journal of Clinical Microbiology & Infectious Diseases 35: 917–925.

Bortolaia, V., Kaas, R.S., Ruppe, E., Roberts, M.C., Schwarz, S., Cattoir, V. et al. (2020) ResFinder 4.0 for predictions of phenotypes from genotypes. Journal of Antimicrobial Chemotherapy 75: 3491–3500.

Bosman, A.L., Loest, D., Carson, C.A., Agunos, A., Collineau, L., and Léger, D.F. (2019) Developing Canadian Defined Daily Doses for Animals: A Metric to Quantify Antimicrobial Use. Frontiers in Veterinary Science 6: 220.

Brenciani, A., Morroni, G., Vincenzi, C., Manso, E., Mingoia, M., Giovanetti, E., and Varaldo, P.E. (2016) Detection in Italy of two clinical *Enterococcus faecium* isolates carrying both the oxazolidinone and phenicol resistance gene optrA and a silent multiresistance gene cfr. Journal of Antimicrobial Chemotherapy 71:1118–1119.

Calland, J.K., Pascoe, B., Bayliss, S.C., Mourkas, E., Berthenet, E., Thorpe, H.A. Hitching, M.D. Feil, E.J. Blaser, M.J. Falush, D. and Sheppard, S.K. (2020) Quantifying bacterial evolution in the wild: a birthday problem for Campylobacter lineages. bioRxiv.

Carattoli, A., Zankari, E., García-Fernández, A., Larsen, M.V., Lund, O., Villa, L. Aarestrup, F.M. and Hasman, H. (2014) In silico detection and typing of plasmids using PlasmidFinder and plasmid multilocus sequence typing. Antimicrobial agents and chemotherapy 58: 3895.

Chen, L., Song, Y., Wei, Z., He, H., Zhang, A., and Jin, M. (2012) Antimicrobial susceptibility, tetracycline and erythromycin resistance genes, and multilocus sequence typing of *Streptococcus suis* Isolates from Diseased Pigs in China. Journal of Veterinary Medical Science 12:0279.

CLSI (2002) Performance standards for antimicrobial disk and dilution susceptibility tests for bacteria isolated from animals; Approved standard-second Edition. In Clinical and Laboratory Standards Institute, Wayne, PA.

CLSI (2008) Performance Standards for Antimicrobial Disk and Dilution Susceptibility Tests for Bacteria Isolated from Animals; Approved Standard. 3rd ed. In Clinical and Laboratory Standards Institute, Wayne, PA.

CLSI (2012) Performance Standards for Antimicrobial Disk Susceptibility Tests; Approved Standard— Eleventh Edition In Clinical and Laboratory Standards Institute, Wayne, PA, p. 76.

CLSI (2017) Performance Standards for Antimicrobial Susceptibility Testing. M100-S27. In Clinical and Laboratory Standards Institute, Wayne, PA.

CLSI (2018) Performance standards for antimicrobial disk and dilution susceptibility tests for bacteria isolated from animals; 4th Edition. In Clinical and Laboratory Standards Institute, Wayne, PA.

CLSI (2020) Performance Standards for Antimicrobial Susceptibility Testing. M100-S30. In Clinical and Laboratory Standards Institute, Wayne, PA.

Coordinators NR (2018) Database resources of the national center for biotechnology information. Nucleic acids research 46: D8–D13.

Dearlove, B.L., Cody, A.J., Pascoe, B., Méric, G., Wilson, D.J., and Sheppard, S.K. (2016) Rapid host switching in generalist Campylobacter strains erodes the signal for tracing human infections. The ISME journal 10: 721–729.

Delannoy, S., Le Devendec, L., Jouy, E., Fach, P., Drider, D., and Kempf, I. (2017) Characterization of Colistin-Resistant *Escherichia coli* Isolated from Diseased Pigs in France. Frontiers in Microbiology 8: 2278.

Du, F., Lv, X., Duan, D., Wang, L., and Huang, J. (2019) Characterization of a linezolid-and vancomycin-resistant *Streptococcus suis* isolate that harbours *optrA* and *vanG* operons. Frontiers in microbiology 10: 2026.

Dutkiewicz, J., Sroka, J., Zając, V., Wasiński, B., Cisak, E., Sawczyn, A. et al. (2017) *Streptococcus suis*: a re-emerging pathogen associated with occupational exposure to pigs or pork products. Part I–Epidemiology. Annals of Agricultural and Environmental Medicine 24: 683–695.

EUCAST- and CLSI potency NEO-SENSITABS™ (2013) Veterinary practice according to CLSI breakpoints.

Flores, J.L., Higgins, R., D’Allaire, S., Charette, R., Boudreau, M., and Gottschalk, M. (1993) Distribution of the different capsular types of *Streptococcus suis* in nineteen swine nurseries. The Canadian veterinary journal 34: 170–171.

Florez-Cuadrado, D., Ugarte-Ruiz, M., Meric, G., Quesada, A., Porrero, M.C., Pascoe, B. Sáez-Llorente, J.L. Orozco, G.L. Domínguez, L. and Sheppard, S.K. (2017) Genome Comparison of Erythromycin Resistant Campylobacter from Turkeys Identifies Hosts and Pathways for Horizontal Spread of *erm*(*B*) Genes. Frontiers in Microbiology 8: 2240.

Gilbert, M., Nicolas, G., Cinardi, G., Van Boeckel, T.P., Vanwambeke, S.O., Wint, G.R.W., and Robinson, T.P. (2018) Global distribution data for cattle, buffaloes, horses, sheep, goats, pigs, chickens and ducks in 2010. Scientific Data 5: 180227.

Gottschalk, M., and Segura, M. (2019) Streptococcosis. Diseases of Swine: 934–950.

Goyette-Desjardins, G., Auger, J.-P., Xu, J., Segura, M., and Gottschalk, M. (2014) *Streptococcus suis*, an important pig pathogen and emerging zoonotic agent: an update on the worldwide distribution based on serotyping and sequence typing. Emerging microbes & infections 3:1–20.

Gurung, M., Tamang, M.D., Moon, D.C., Kim, S.-R., Jeong, J.-H., Jang, G.-C. et al. (2015) Molecular basis of resistance to selected antimicrobial agents in the emerging zoonotic pathogen *Streptococcus suis*. Journal of clinical microbiology 53: 2332–2336.

Hadjirin, N.F., Miller, E.L., Murray, G.G.R., Yen, P.L.K., Phuc, H.D., Wileman, T.M. et al. (2020) Linking phenotype, genotype and ecology: antimicrobial resistance in the zoonotic pathogen *Streptococcus suis*. bioRxiv.

Hadjirin, N.F., Miller, E.L., Murray, G.G.R., Yen, P.L.K., Phuc, H.D., Wileman, T.M. et al. (2021) A comprehensive portrait of antimicrobial resistance in the zoonotic pathogen *Streptococcus suis*. bioRxiv.

Hatrongjit, R., Fittipaldi, N., Gottschalk, M., and Kerdsin, A. (2020) Tools for Molecular Epidemiology of *Streptococcus suis*. Pathogens 9.2: 81.

Hoa, N.T., Chieu, T.T.B., Nghia, H.D.T., Mai, N.T.H., Anh, P.H., Wolbers, M. et al. (2011) The antimicrobial resistance patterns and associated determinants in *Streptococcus suis* isolated from humans in southern Vietnam, 1997-2008. BMC infectious diseases 11: 6–6.

Holden, M.T., Hauser, H., Sanders, M., Ngo, T.H., Cherevach, I., Cronin, A. et al. (2009) Rapid evolution of virulence and drug resistance in the emerging zoonotic pathogen *Streptococcus suis*. PloS one 4:7, e6072.

Howe, R.A., and Andrews, J.M. (2012) BSAC standardized disc susceptibility testing method (version 11). Journal of antimicrobial chemotherapy 67: 2783–2784.

Huang, J., Chen, L., Wu, Z., and Wang, L. (2017) Retrospective analysis of genome sequences revealed the wide dissemination of optrA in Gram-positive bacteria. Journal of Antimicrobial Chemotherapy 72: 614–616.

Huang, J., Sun, J., Wu, Y., Chen, L., Duan, D., Lv, X., and Wang, L. (2019) Identification and pathogenicity of an XDR *Streptococcus suis* isolate that harbours the phenicol-oxazolidinone resistance genes *optrA* and *cfr*, and the bacitracin resistance locus *bcrABDR*. International journal of antimicrobial agents 54: 43–48.

Huang, K., Zhang, Q., Song, Y., Zhang, Z., Zhang, A., Xiao, J., and Jin, M. (2016) Characterization of spectinomycin resistance in *Streptococcus suis* leads to two novel insights into drug resistance formation and dissemination mechanism. Antimicrobial agents and chemotherapy 60: 6390–6392.

Hughes, J.M., Wilson, M.E., Wertheim, H.F., Nghia, H.D.T., Taylor, W., and Schultsz, C. (2009) *Streptococcus suis*: an emerging human pathogen. Clinical Infectious Diseases 48: 617–625.

Katoh, K., Misawa, K., Kuma, K.i., and Miyata, T. (2002) MAFFT: a novel method for rapid multiple sequence alignment based on fast Fourier transform. Nucleic Acids Research 30: 3059–3066.

Kerdsin, A., Hatrongjit, R., Gottschalk, M., Takeuchi, D., Hamada, S., Akeda, Y., and Oishi, K. (2017) Emergence of *Streptococcus suis* serotype 9 infection in humans. Journal of microbiology, immunology, and infection= Wei mian yu gan ran za zhi 50: 545–546.

Kongkaew, S., Wongsawan, K., Pansumdang, C., Takam, S., Yano, T., Yamsakul, P., and Patchanee, P. (2012) Identification and antimicrobial susceptibility of *Streptococcus suis* isolated from pigs tonsil swabs. Kasetsart Veterinarians 22: 1–13.

Lakkitjaroen, N., Kaewmongkol, S., Metheenukul, P., Karnchanabanthoeng, A., Satchasataporn, K., Abking, N., and Rerkamnuaychoke, W. (2011) Prevalence and antimicrobial susceptibility of *Streptococcus suis* isolated from slaughter pigs in Northern Thailand. Kasetsart J (Nat Sci) 45: 78–83.

Lees, J.A., Harris, S.R., Tonkin-Hill, G., Gladstone, R.A., Lo, S.W., Weiser, J.N. et al. (2019) Fast and flexible bacterial genomic epidemiology with PopPUNK. Genome research 29: 304–316.

Li, X.-S., Dong, W.-C., Wang, X.-M., Hu, G.-Z., Wang, Y.-B., Cai, B.-Y. et al. (2014) Presence and genetic environment of pleuromutilin–lincosamide–streptogramin A resistance gene *lsa* (*E*) in enterococci of human and swine origin. Journal of Antimicrobial Chemotherapy 69: 1424–1426.

Liu, B., Zheng, D., Jin, Q., Chen, L., and Yang, J. (2019) VFDB 2019: a comparative pathogenomic platform with an interactive web interface. Nucleic Acids Research 47: D687–D692.

Liu, Y.-Y., Wang, Y., Walsh, T.R., Yi, L.-X., Zhang, R., Spencer, J. et al. (2016) Emergence of plasmid-mediated colistin resistance mechanism MCR-1 in animals and human beings in China: a microbiological and molecular biological study. The Lancet Infectious Diseases 16: 161–168.

Maisey, H.C., Hensler, M., Nizet, V., and Doran, K.S. (2007) Group B Streptococcal Pilus Proteins Contribute to Adherence to and Invasion of Brain Microvascular Endothelial Cells. Journal of Bacteriology 189: 1464.

Marois, C., Bougeard, S., Gottschalk, M., and Kobisch, M. (2004) Multiplex PCR assay for detection of *Streptococcus suis* species and serotypes 2 and 1/2 in tonsils of live and dead pigs. Journal of clinical microbiology 42: 3169–3175.

Martel, A., Baele, M., Devriese, L., Goossens, H., Wisselink, H., Decostere, A., and Haesebrouck, F. (2001) Prevalence and mechanism of resistance against macrolides and lincosamides in *Streptococcus suis* isolates. Veterinary microbiology 83: 287–297.

McHugh, M.P., Parcell, B.J., Pettigrew, K.A., Toner, G., Khatamzas, E., Karcher, A.M. et al. (2020) Emergence of *optrA*-mediated linezolid resistance in multiple lineages and plasmids of Enterococcus faecalis revealed by long read sequencing. bioRxiv.

McInerney, J.O., McNally, A., and O’Connell, M.J. (2017) Why prokaryotes have pangenomes. Nature Microbiology 2: 17040.

Morley, V.J., Woods, R.J., and Read, A.F. (2019) Bystander selection for antimicrobial resistance: implications for patient health. Trends in microbiology 27: 864–877.

Mourkas, E., Florez-Cuadrado, D., Pascoe, B., Calland, J.K., Bayliss, S.C., Mageiros, L. et al. (2019) Gene pool transmission of multi-drug resistance among Campylobacter from livestock, sewage and human disease. Environmental Microbiology 21: 4597–4613.

Mourkas, E., Taylor, A.J., Méric, G., Bayliss, S.C., Pascoe, B., Mageiros, L. et al. (2020) Agricultural intensification and the evolution of host specialism in the enteric pathogen *Campylobacter jejuni*. Proceedings of the National Academy of Sciences 117: 11018.

Murray, G.G.R., Charlesworth, J., Miller, E.L., Casey, M.J., Lloyd, C.T., Gottschalk, M. et al. (2021) Genome reduction is associated with bacterial pathogenicity across different scales of temporal and ecological divergence. Molecular Biology and Evolution 38: 1570–1579.

National Research Council (2010) Guide for the care and use of laboratory animals: National Academies Press.

Nguyen, L.-T., Schmidt, H.A., Von Haeseler, A., and Minh, B.Q. (2015) IQ-TREE: a fast and effective stochastic algorithm for estimating maximum-likelihood phylogenies. Molecular biology and evolution 32: 268–274.

Nguyen, N.T., Nguyen, H.M., Nguyen, C.V., Nguyen, T.V., Nguyen, M.T., Thai, H.Q. et al. (2016) Use of colistin and other critical antimicrobials on pig and chicken farms in Southern Vietnam and its association with resistance in commensal *Escherichia coli*. bacteria. Applied and Environmental Microbiology 82: 3727.

Okura, M., Osaki, M., Nomoto, R., Arai, S., Osawa, R., Sekizaki, T., and Takamatsu, D. (2016) Current taxonomical situation of *Streptococcus suis*. Pathogens 5: 45.

Okura, M., Maruyama, F., Ota, A., Tanaka, T., Matoba, Y., Osawa, A. et al. (2019) Genotypic diversity of *Streptococcus suis* and the *S. *suis*-like bacterium *Streptococcus ruminantium* in ruminants*. Veterinary Research 50: 94.

Padungtod, P., Tharavichitkul, P., Junya, S., Chaisowong, W., Kadohira, M., Makino, S., and Sthitmatee, N. (2010) Incidence and presence of virulence factors of *Streptococcus suis* infection in slaughtered pigs from Chiang Mai, Thailand. Southeast Asian journal of tropical medicine and public health 41: 1454.

Palmieri, C., Varaldo, P.E., and Facinelli, B. (2011) *Streptococcus suis*, an emerging drug-resistant animal and human pathogen. Frontiers in microbiology 2: 235.

Patchanee, P., Chokesajjawatee, N., Santiyanont, P., Chuammitri, P., Deeudom, M., Monteith, W. et al. (2020a) Multiple clones of colistin-resistant Salmonella enterica carrying plasmids in meat products and patients in Northern Thailand. bioRxiv.

Patchanee, P., Tanamai, P., Tadee, P., Hitchings, M.D., Calland, J.K., Sheppard, S.K. et al. (2020b) Whole-genome characterisation of multi-drug resistant monophasic variants of Salmonella Typhimurium from pig production in Thailand. PeerJ 8: e9700.

Pathanasophon, P., Worarach, A., Narongsak, W., Yuwapanichsampan, S., Nuangmek, A., Sakdasirisathaporn, A., and Chuxnum, T. (2013) Prevalence of *Streptococcus suis* in Tonsils of Slaughtered Pigs in Lampang and Phayao Provinces, Thailand, 2009-2010. Journal of Tropical Medicine & Parasitology 36: 8–14.

Prasertsee, T., Chuammitri, P., Deeudom, M., Chokesajjawatee, N., Santiyanont, P., Tadee, P. et al. (2019) Core genome sequence analysis to characterize *Salmonella enterica* serovar Rissen ST469 from a swine production chain. International journal of food microbiology 304: 68–74.

Prüfer, T.L., Rohde, J., Verspohl, J., Rohde, M., De Greeff, A., Willenborg, J., and Valentin-Weigand, P. (2019) Molecular typing of *Streptococcus suis* strains isolated from diseased and healthy pigs between 1996-2016. PloS one 14: e0210801.

Pumart P, P.T., Thamlikitkul V, Riewpaiboon A, Prakongsai P, Limwattananon S (2012) Health and economic impacts of antimicrobial resistance in Thailand. J Health Serv Res Policy 6: 352–360.

Quinn, P.J., Carter, M.E., Markey, B., and Carter, G.R. (1994) Clinical Veterinary Microbiology: Wolfe.

Rayanakorn, A., Ademi, Z., Liew, D., and Lee, L.H. (2020) PIN65 Estimating the lifetime economic burden of *Streptococcus suis* and its productivity impact in Thailand. Value in Health 23: S179.

Rayanakorn, A., Katip, W., Goh, B.H., Oberdorfer, P., and Lee, L.H. (2019) Clinical Manifestations and Risk Factors of *Streptococcus suis* Mortality Among Northern Thai Population: Retrospective 13-Year Cohort Study. Infect Drug Resist 12: 3955–3965.

Redondo-Salvo, S., Fernández-López, R., Ruiz, R., Vielva, L., de Toro, M., Rocha, E.P.C. et al. (2020) Pathways for horizontal gene transfer in bacteria revealed by a global map of their plasmids. Nature Communications 11: 3602.

Robertson, J., and Nash, J.H.E. (2018) MOB-suite: software tools for clustering, reconstruction and typing of plasmids from draft assemblies. Microbial genomics 4:8.

Robertson, J., Bessonov, K., Schonfeld, J., and Nash, J.H.E. (2020) Universal whole-sequence-based plasmid typing and its utility to prediction of host range and epidemiological surveillance. Microbial Genomics 6:10.

Seemann, T. (2014) Prokka: rapid prokaryotic genome annotation. Bioinformatics 30: 2068–2069.

Segura, M. (2020) *Streptococcus suis* Research: Progress and Challenges. Pathogens 9:707.

Segura, M., Calzas, C., Grenier, D., and Gottschalk, M. (2016) Initial steps of the pathogenesis of the infection caused by *Streptococcus suis*: fighting against nonspecific defenses. FEBS letters 590: 3772–3799.

Segura, M., Aragon, V., Brockmeier, S.L., Gebhart, C., Greeff, A.d., Kerdsin, A. et al. (2020) Update on *Streptococcus suis* Research and Prevention in the Era of Antimicrobial Restriction: 4th International Workshop on S. suis. Pathogens (Basel, Switzerland) 9: 374.

Seitz, M., Valentin-Weigand, P., and Willenborg, J. (2016) Use of antibiotics and antimicrobial resistance in veterinary medicine as exemplified by the swine pathogen *Streptococcus suis*. In How to Overcome the Antibiotic Crisis: Springer, pp. 103–121.

Sheppard, S.K., Guttman, D.S., and Fitzgerald, J.R. (2018) Population genomics of bacterial host adaptation. Nature Reviews Genetics 19: 549–565.

Soares, T.C.S., Paes, A.C., Megid, J., Ribolla, P.E.M., Paduan, K.d.S., and Gottschalk, M. (2014) Antimicrobial susceptibility of *Streptococcus suis* isolated from clinically healthy swine in Brazil. Canadian journal of veterinary research 78: 145–149.

Stevens, M.J.A., Spoerry Serrano, N., Cernela, N., Schmitt, S., Schrenzel, J., and Stephan, R. (2019) Massive diversity in whole-genome sequences of *Streptococcus suis* strains from infected pigs in Switzerland. Microbiology Resource Announcements 8.5: e01656–18.

Sweeney, M.T., Lubbers, B.V., Schwarz, S., and Watts, J.L. (2018) Applying definitions for multi-drug resistance, extensive drug resistance and pandrug resistance to clinically significant livestock and companion animal bacterial pathogens. Journal of Antimicrobial Chemotherapy 73: 1460–1463.

Sztanke, K., Pasternak, K., and Sztanke, M. (2004) Oxazolidinones--a new class of broad-spectrum chemotherapeutics. In Annales Universitatis Mariae Curie-Sklodowska Sectio D: Medicina, pp. 335–341.

Tadee, P., Patchanee, P., Pascoe, B., Sheppard, S.K., Meunsene, D., Buawiratlert, T., and Tadee, P. (2021) Occurrence and sequence type of antimicrobial resistant *Salmonella spp*. circulating in antibiotic-free organic pig farms of northern-Thailand. The Thai Journal of Veterinary Medicine 51: 311–319.

Takeuchi, D., Kerdsin, A., Akeda, Y., Chiranairadul, P., Loetthong, P., Tanburawong, N. et al. (2017) Impact of a food safety campaign on *Streptococcus suis* infection in humans in Thailand. The American journal of tropical medicine and hygiene 96: 1370–1377.

Tan, M.F., Tan, J., Zeng, Y.B., Li, H.Q., Yang, Q., and Zhou, R. (2020) Antimicrobial resistance phenotypes and genotypes of *Streptococcus suis* isolated from clinically healthy pigs from 2017 to 2019 in Jiangxi Province, China. Journal of Applied Microbiology 130.3: 797–806.

Tedijanto, C., Olesen, S.W., Grad, Y.H., and Lipsitch, M. (2018) Estimating the proportion of bystander selection for antibiotic resistance among potentially pathogenic bacterial flora. Proceedings of the National Academy of Sciences 115: e11988–95.

Thongkamkoon, P., Kiatyingangsulee, T., and Gottschalk, M. (2017) Serotypes of *Streptococcus suis* isolated from healthy pigs in Phayao Province, Thailand. BMC research notes 10: 53.

Van Boeckel, T.P., Brower, C., Gilbert, M., Grenfell, B.T., Levin, S.A., Robinson, T.P. et al. (2015) Global trends in antimicrobial use in food animals. Proceedings of the National Academy of Sciences 112: 5649–5654.

van Samkar, A., Brouwer, M.C., Schultsz, C., van der Ende, A., and van de Beek, D. (2015) *Streptococcus suis* meningitis: a systematic review and meta-analysis. PLoS Negl Trop Dis 9: e0004191.

VanderWaal, K., and Deen, J. (2018) Global trends in infectious diseases of swine. Proceedings of the National Academy of Sciences 115: 11495.

Wang, B., Wang, Y., Xie, X., Diao, Z., Xie, K., Zhang, G. et al. (2020) Quantitative Analysis of Spectinomycin and Lincomycin in Poultry Eggs by Accelerated Solvent Extraction Coupled with Gas Chromatography Tandem Mass Spectrometry. Foods 9: 651.

Wang, R., van Dorp, L., Shaw, L.P., Bradley, P., Wang, Q., Wang, X. et al. (2018) The global distribution and spread of the mobilized colistin resistance gene *mcr-1*. Nature Communications 9: 1179.

Wang, Y., Lv, Y., Cai, J., Schwarz, S., Cui, L., Hu, Z. et al. (2015) A novel gene, *optrA*, that confers transferable resistance to oxazolidinones and phenicols and its presence in *Enterococcus faecalis* and *Enterococcus faecium* of human and animal origin. Journal of Antimicrobial Chemotherapy 70: 2182–2190.

Weinert, L.A., and Welch, J.J. (2017) Why Might bacterial pathogens have small genomes?. Trends in Ecology & Evolution 32: 936–947.

Weinert, L.A., Chaudhuri, R.R., Wang, J., Peters, S.E., Corander, J., Jombart, T. et al. (2019) Publisher Correction: Genomic signatures of human and animal disease in the zoonotic pathogen Streptococcus suis. Nature communications 10: 5326–5326.

Weinert, L.A., Chaudhuri, R.R., Wang, J., Peters, S.E., Corander, J., Jombart, T. et al. (2015) Genomic signatures of human and animal disease in the zoonotic pathogen *Streptococcus suis*. Nature Communications 6: 6740.

World Health Organization (2017) WHO guidelines on use of medically important antimicrobials in food-producing animals: web annex A: evidence base. In: World Health Organization.

Wisselink, H.J., Joosten, J.J., and Smith, H.E. (2002) Multiplex PCR Assays for Simultaneous Detection of Six Major Serotypes and Two Virulence-Associated Phenotypes of *Streptococcus suis* in Tonsillar Specimens from Pigs. Journal of Clinical Microbiology 40: 2922–2929.

Wisselink, H.J., Smith, H.E., Stockhofe-Zurwieden, N., Peperkamp, K., and Vecht, U. (2000) Distribution of capsular types and production of muramidase-released protein (MRP) and extracellular factor (EF) of *Streptococcus suis* strains isolated from diseased pigs in seven European countries. Veterinary microbiology 74: 237–248.

Yan, H., Yu, R., Li, D., Shi, L., Schwarz, S., Yao, H. et al. (2020) A novel multiresistance gene cluster located on a plasmid-borne transposon in *Listeria monocytogenes*. Journal of Antimicrobial Chemotherapy 75: 868–872.

Yongkiettrakul, S., Maneerat, K., Arechanajan, B., Malila, Y., Srimanote, P., Gottschalk, M., and Visessanguan, W. (2019) Antimicrobial susceptibility of *Streptococcus suis* isolated from diseased pigs, asymptomatic pigs, and human patients in Thailand. BMC veterinary research 15: 5.

Young, J.P.W. (2016) Bacteria are smartphones and mobile genes are apps. Trends in microbiology 24: 931–932.

Zhang, A., Yang, M., Hu, P., Wu, J., Chen, B., Hua, Y. et al. (2011) Comparative genomic analysis of *Streptococcus suis* reveals significant genomic diversity among different serotypes. BMC Genomics 12: 523.

Zhang, C., Zhang, Z., Song, L., Fan, X., Wen, F., Xu, S., and Ning, Y.J.B.r.i. (2015) Antimicrobial resistance profile and genotypic characteristics of *Streptococcus suis* capsular type 2 isolated from clinical carrier sows and diseased pigs in China. BioMed research international 2015.

Zhao, S., Tyson, G.H., Chen, Y., Li, C., Mukherjee, S., Young, S. et al. (2016) Whole-genome sequencing analysis accurately predicts antimicrobial resistance phenotypes in *Campylobacter spp*. Applied and Environmental Microbiology 82: 459–466.

Zhou, W., Gao, S., Xu, H., Zhang, Z., Chen, F., Shen, H., and Zhang, C. (2019) Distribution of the *optrA* gene in Enterococcus isolates at a tertiary care hospital in China. Journal of global antimicrobial resistance 17: 180–186.

