## Supplementary figures and images for "Non-serotype 2 isolates from healthy pigs are a potential zoonotic reservoir of *Streptococcus suis* genetic diversity and antimicrobial resistance"

### Figure S1

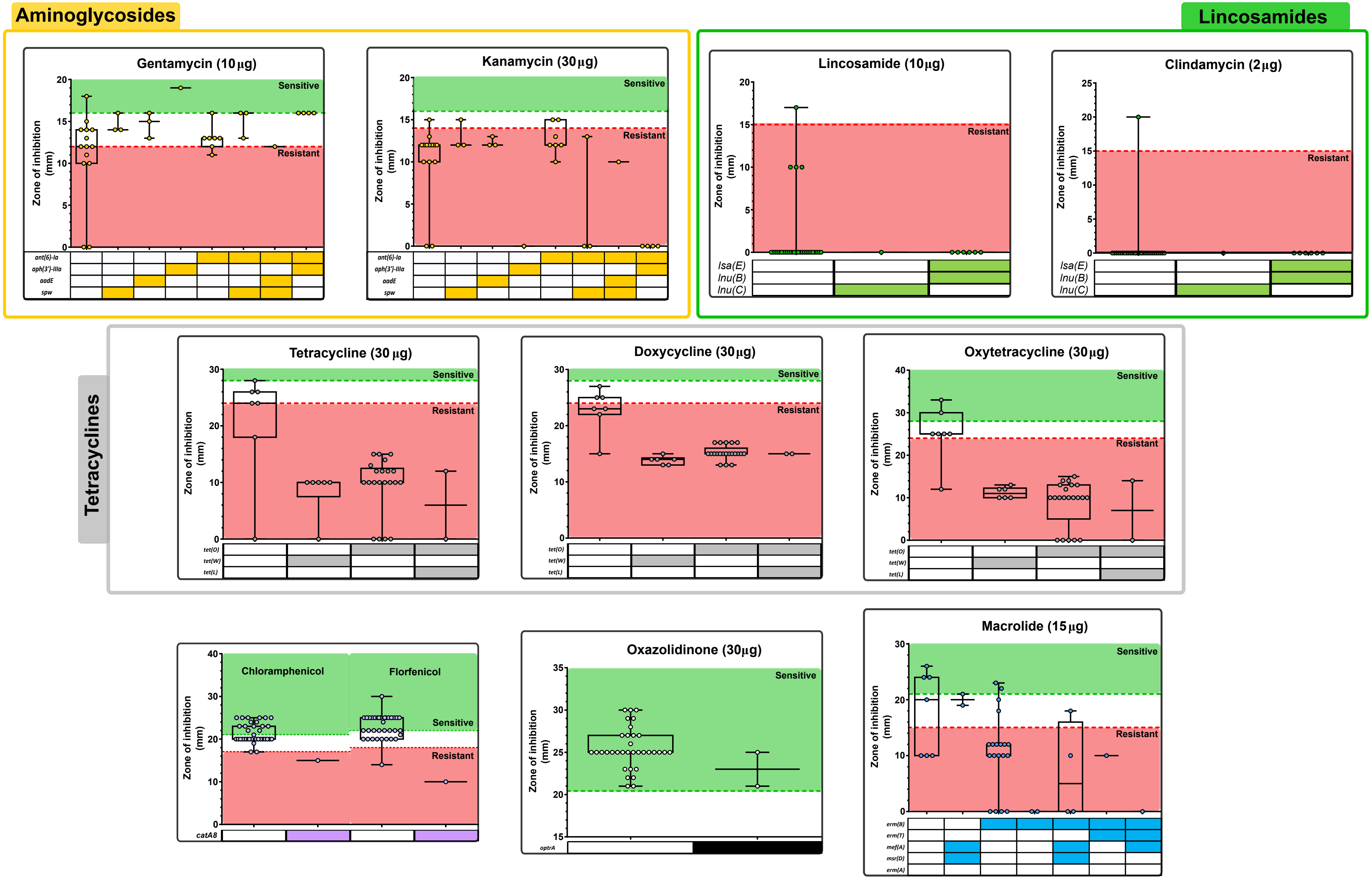
